# RAINSTORM: Automated Behavioral Analysis for Mice Exploratory Behavior Using Artificial Neural Networks

**DOI:** 10.1101/2025.04.07.647548

**Authors:** Santiago D’hers, Agustina Denise Robles, Santiago Ojea Ramos, Guillermina Bollini, Mariana Feld

## Abstract

Rodent exploratory behavior is widely used for assessing cognitive function. RAINSTORM — Real and Artificial Intelligence for Neuroscience - Simple Tracker for Object Recognition Memory — is a versatile tool designed to streamline the analysis of such behaviors in rodents. This pipeline integrates manual behavioral scoring, geometric analysis, and artificial intelligence (AI)-powered behavioral labeling, offering reproducible, scalable, and efficient evaluation methods. RAINSTORM processes raw positional data and automates the identification of exploratory behaviors, providing insights into memory performance.

This tool is designed to learn from the labeling criteria of one or more experimenters by capturing the different aspects of expert opinion and reducing subjective bias in subsequent scoring procedures. The experimenter can go from unprocessed pose estimation data (obtained through open source software such as DeepLabCut) to accurate exploration patterns in a matter of minutes. By optimizing the analysis process, RAINSTORM significantly enhances the reliability and efficiency of behavioral research.

RAINSTORM has become a robust methodology for assessing recognition memory in rodents by accurately quantifying exploration times for familiar and novel objects. It has since been extended to include (but not limited to) a wider range of exploratory behaviors. The software is readily applicable to other experimental designs that rely on quantifying exploration in mice, such as Social Preference and Object Pattern Separation, among others.

## Introduction

Manual scoring of animal behavior, particularly in studies involving exploration, is not only time-consuming but also imprecise and susceptible to operator bias. It relies on the expertise of few researchers, which reduces reproducibility of results (Tuyttens et al., 2014). On the other hand, commercial tracking systems have become popular for animal behavior analysis due to their user-friendly interfaces and integrated experimental management (Antunes et al., 2011; Noldus et al., 2001). However, these systems typically employ **heuristic methods** that oversimplify exploratory behavior to basic metrics like the distance and angle of approach, and tend to underperform compared to both manual scoring and advanced **machine learning** techniques. These tools lack transparency, as users cannot examine or modify the underlying algorithms, making it difficult to search for accuracy improvements or adapt them to specific research needs. Additionally, their high cost further limits accessibility.

Recently, there has been a surge in **open-source tools** that empower non-experts to process experimental data, and provide a flexible solution for the analysis of behavioral recordings (Pennington et al., 2021; Rodriguez et al., 2018). However, their reliance on fixed, heuristic-based algorithms limits their ability to capture the nuanced spectrum of exploratory actions. Recognizing these limitations, a new wave of collaborative open-access software has emerged that integrates machine learning techniques, enabling models to dynamically learn and adapt to the subtle complexities of exploratory behavior. These advancements represent a significant leap from traditional heuristic methods toward data-driven approaches, ultimately delivering a more complete and automated analysis pipeline integrating the flexibility needed to capture experimenter-defined behavioral criteria.

One of the most widely used tools that incorporate machine learning to video analysis is **DeepLabCut** (DLC, Mathis et al., 2018), a toolbox for state-of-the-art markerless pose estimation of animals performing diverse behaviors. Pose estimation software processes videos where rodents (or other animals) display their behaviors, and return the tracked coordinates for any number of selected body parts. These tools are very useful in automating the analysis of the behavior, but they may leave the experimenter quite a bit far behind the final result. Although some tools have been developed for processing positions for the analysis of behaviors (Isik & Unal, 2023), none to our knowledge covers all the steps from video acquisition to visualization of results. These tools greatly facilitate the automation of video processing, but often fall short of delivering a complete analysis process—from video acquisition to the visualization of behavioral patterns. They either fail to capture the experimenter’s criteria to identify exploratory behaviors, or the computer setup becomes quite complex in the search for a more generalized tool for more complicated behaviors.

To address these challenges, we have developed **RAINSTORM**, a novel automated behavioral analysis method that uses Python-based **Artificial Neural Networks** (ANN). Building on advances in frameworks like TensorFlow (Abadi et al., 2016), which have simplified the development and training of deep learning models in Python, RAINSTORM goes beyond the limitations of earlier heuristic approaches. This tool makes use of the pose estimation data obtained with pose estimation software (i.e. DLC) to train automated Artificial Intelligence (AI) models for behavior analysis. Following this protocol, users will obtain detailed quantification of mouse behaviors in a fast, reliable, reusable, and straightforward manner. As an open-source tool, RAINSTORM is freely available, allowing users to access the backend, modify its functionality, and tailor it to their specific research goals.

Users can manually label videos frame-by-frame with RAINSTORM’s Behavioral Labeler tool, analyze animal positions through geometric behavioral labeling, or leverage the power of AI training, evaluation and automated behavioral labeling. Beginners and non-programmers can train a custom artificial neural network using their own labeled data, allowing them to categorize new videos according to expert-defined criteria.

### Strategic Planning

RAINSTORM offers three distinct methods for behavioral labeling: Manual, Geometric, and Automated, along with support protocols for installing the package and running video processing for better pose estimation performance (Fig. 1). The **Manual** labeling method allows users to annotate each video frame directly within the RAINSTORM Behavioral Labeler by assigning behavioral categories, event markers, or other relevant labels based on their personal observations.

**Figure 1.**
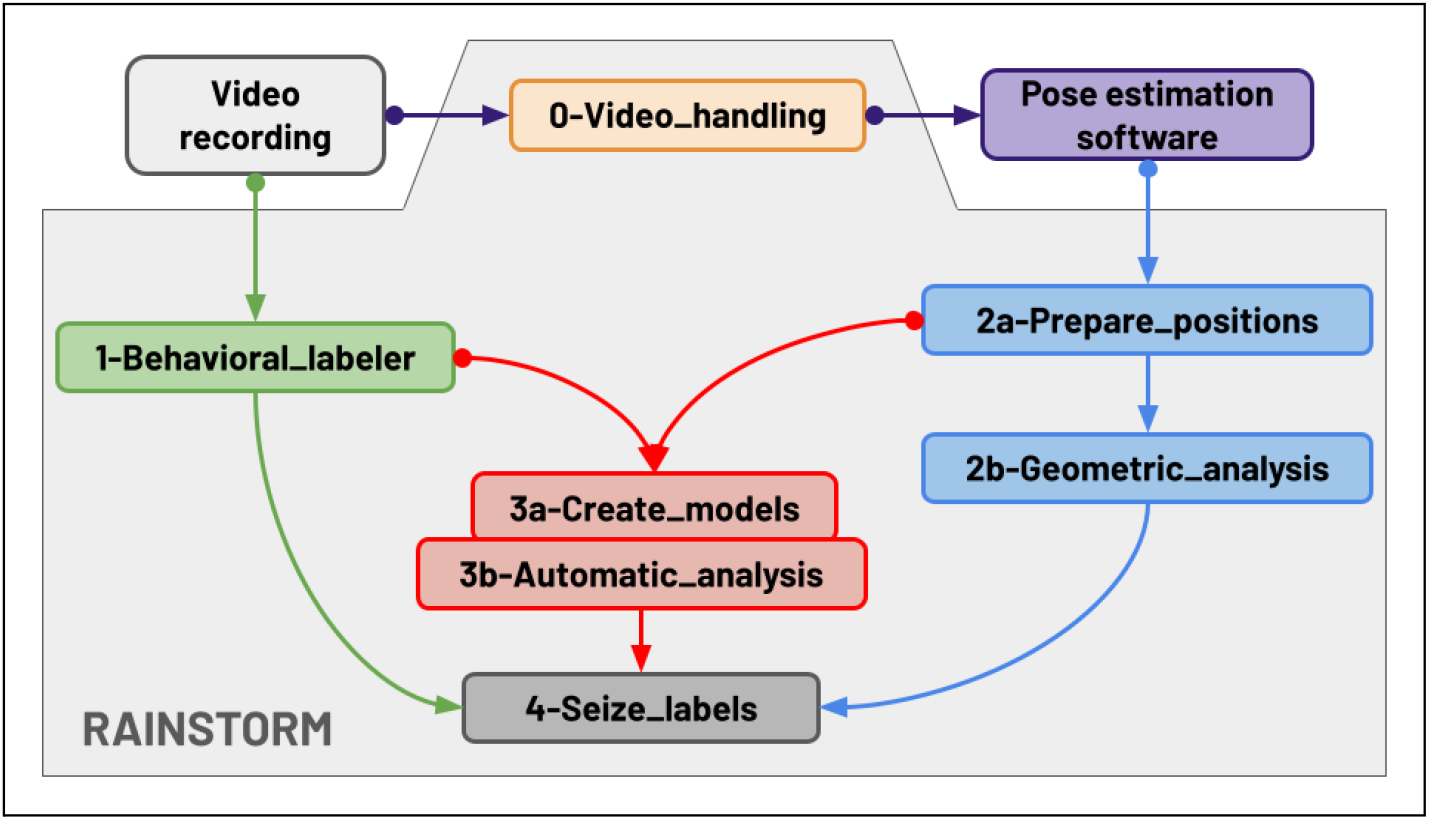
Workflow diagram. Video recordings (light gray box) can be analyzed with the RAINSTORM **1-Behavioral_labeler** (green box), or can be prepared using **0-Video_handling** (orange box) to be processed with a pose estimation software (e.g. DeepLabCut or SLEAP, purple box) and analyzed with the geometric method (**2a-Prepare_positions** and **2b-Geometric_analysis**, blue boxes). Labels and positions can be used to train an Artificial Neural Network (**3a-Create_models** and **3b-Automatic_analysis**, red boxes). Any of the three paths lead to the visualization of the final results in **4-Seize_labels** (dark gray box).

In contrast, the **Geometric** and **Automatic** methods utilize mouse position data obtained through pose estimation software (e.g., DLC or SLEAP) but differ in their analytical approaches. The Geometric method classifies animal behavior arithmetically, using its position, general movement and user defined thresholds. It assesses exploratory behavior by analyzing the positions of the mouse’s head and nose relative to the target, using metrics such as distance and angle of approach. The Automated method leverages both mouse position data and manually labeled behaviors to train artificial neural networks, integrating experimenter-defined criteria to enable the automated labeling of exploratory behaviors. Together, these methods provide a versatile toolkit for behavioral analysis, addressing varying levels of detail and automation.

### Support Protocol 1

RAINSTORM Package Installation

#### Introductory paragraph

RAINSTORM is an open source tool built entirely with free and widely used software. It integrates with platforms such as **Anaconda, Git** and **Visual Studio Code**, and ensures accessibility and ease of use for researchers and developers. The package is hosted on the Python Package Index (PyPI), so it can be easily installed by running ‘pip install rainstorm’. This section will guide you through the steps required to set up RAINSTORM and get started.

#### Necessary Resources

The software was fully developed and tested on a notebook with i5 processor and 8 GB of RAM, with no GPU. Although it does not need sophisticated computer hardware, for a better experience it is recommended to use a computer with at least 16 GB RAM (since some parts of the processing can be demanding for the computer memory).

For users that would like to train the Neural Networks (Fig. 1, step 3a-Create_models) with GPU acceleration but lack access to a local GPU, Google Colab offers a free and efficient alternative to harness the power of cloud-based GPUs.

#### Protocol steps with *step annotations*

1. Install Miniconda (or Anaconda), Visual Studio Code and Git.
2. *Miniconda: https://docs.anaconda.com/miniconda/install* *During the installation process, select “Add Miniconda3 to my path”*.
3. *Visual Studio Code: https://code.visualstudio.com/download*
4. *Git: https://git-scm.com/downloads*

*Having Git installed is optional, but recommended, since you can download the RAINSTORM GitHub repository manually from https://github.com/sdhers/RAINSTORM instead of cloning it*.

*Once the three components are installed, it is recommended to restart your computer to ensure that all configurations are properly applied and the system is fully operational*.

5. Clone the RAINSTORM repository.

*Open a Terminal (e.g. Miniconda or Anaconda Prompt). You can see the current working directory path displayed after the environment name:*

~~~
(base) C:\Users\YourUsername>
~~~

*To clone the repository, you will run the following command (This step will create a folder named RAINSTORM on your working directory):*

~~~
(base) C:\Users\YourUsername> git clone https://github.com/sdhers/rainstorm.git
~~~

6. Set up the virtual environment.

*Navigate to the RAINSTORM directory:*

~~~
(base) C:\Users\YourUsername> cd rainstorm
~~~

*Create the Conda Environment:*

~~~
(base) C:\Users\YourUsername> conda env create -f rainstorm_venv.yml
~~~

*This step can take a while (depending on your internet connection). Once the environment is ready, you can activate it by running the following command:*

~~~
(base) C:\Users\YourUsername> conda activate rainstorm (rainstorm) C:\Users\YourUsername>
~~~

7. Open VS code.

*You can launch VS code from the Terminal using:*

~~~
(rainstorm) C:\Users\YourUsername> code .
~~~

*Once in VS code, you need to ensure the Python extension is installed:*

- *Go to the Extensions view (*`Ctrl+Shift+X` *or* `Cmd+Shift+X` *on macOS)*.
- *Search for ‘Python’ and ‘Jupyter’ extensions and install them*.

7. Open the 0-Video_handling Jupyter Notebook.

*Each time you run the first cell of a Notebook, you will be prompted to select a kernel (visible at the top center of the VS Code window). From the available Python environments, choose the RAINSTORM Conda environment. Run the cells in the Notebook to get started. From this point on, you can also launch any of the Notebooks without opening the Terminal (see Table 1 for definitions of computing terminology.) The initial setup is now complete*.

### Support Protocol 2

Video handling

**Table 1.** Glossary of Essential Terms for Interactive and Collaborative Computing.

|  |  |
| --- | --- |
| *GitHub Repository* | *A storage space on GitHub used to organize, store, and manage project files, including code, documentation, and other resources.* |
| *Terminal* | *A text-based interface that allows users to interact with their computer's operating system by typing commands.* |
| *Conda Environment* | *An isolated environment created by the Conda package manager that encapsulates its own set of packages and dependencies.* |
| *Working Directory* | *The folder or directory on your computer where your project files are stored and where the execution of programs or scripts takes place.* |
| *Jupyter Notebook* | *A web-based interactive computing environment where you can combine live code, equations, visualizations, and narrative text in a single document.* |
| *Kernel* | *The computational engine in a Jupyter Notebook that executes the code contained in its cells.* |
| *Cell* | *A container within a Jupyter Notebook that holds code, markdown (text), or other content. Each cell can be executed independently.* |
| *Graphical User Interface (GUI)* | *Interface that allows users to interact with electronic devices through graphical icons and visual indicators.* |

#### Introductory paragraph

This support protocol provides a comprehensive, step-by-step guide for handling and editing video files using a custom Jupyter Notebook with integrated RAINSTORM functions. The protocol is designed to conduct the video editing process, allowing users to efficiently select, trim, crop, align, and annotate video data for subsequent behavioral analysis. By following these standardized procedures, researchers can ensure consistency and accuracy in video preprocessing, thus facilitating robust data analysis and reproducible results.

#### Necessary Resources

After completing the Installation, the necessary **RAINSTORM conda environment** is already set up. Visual Studio Code is recommended for easy navigation through the downloaded repository, allowing users to quickly locate and manage Notebooks and other files. It is highly recommended to use a computer mouse, since some tools require the use of the mouse wheel.

Users may try out the video handling tools on the demo data provided in the downloaded repository; inside `‘docs\examples\NOR\TS_videos’`. The ROIs.json file found in the experiment folder was created using the draw_rois() function found at the end of this support protocol.

#### Protocol steps with *step annotations*

1. Open the Jupyter Notebook: 0-Video_handling.ipynb.

*On VS code, you can navigate through your computer and find the Notebooks in the downloaded repository folder using the Explorer (Ctrl + Shift + E)*.

*Run the first cell to load the necessary modules, including RAINSTORM*.

2. Create a ‘video dictionary’ where you can store the parameters to edit each video.

*Select the videos you want to edit from the pop up window*.

~~~
video_dict = rst.create_video_dict()
~~~

3. Select the time you want the video to start and end.

*Select the start and end time on the pop up window*.

~~~
rst.select_trimming(video_dict)
~~~

4. Select the area of the video you want to crop.

*Draw a rectangular area on the displayed image (built merging frames from all the selected videos)*.

~~~
rst.select_cropping(video_dict)
~~~

*How to use:*

- *Left-click and drag to draw a rectangle*
  ○ *Right-click and drag to move the rectangle*
  ○ *Use the scroll wheel to resize the rectangle*
  ○ *Use Ctrl + scroll wheel to rotate the rectangle*
- *Press ‘c’ to confirm the cropping area*
- *Press ‘q’ to quit*

5. Select the same two points on each video to align them together.

*Click to select two aligning points for each video frame shown*.

~~~
rst.select_alignment(video_dict)
~~~

*How to use:*

- *Left-click to select a point*
- *Press enter to confirm*
- *Repeat until two points are selected for each video*

*It is important to select the same ‘first’ and ‘second’ points each time, since the function will align them considering the selection order*.

6. Apply all the selected transformations to the videos

*The following function will create a folder and store the modified videos*

~~~
rst.apply_transformations(video_dict, trim = True, crop = True, align = True)
~~~

7. Draw Regions of Interest (ROIs)

*This last function allows drawing areas and points on the video, and stores them to use later for behavioral analysis*.

~~~
rst.draw_rois()
~~~

*How to use:*

- *Select the videos you want to draw on. The merged frame will appear (Fig. 2):*

**Figure 2.**
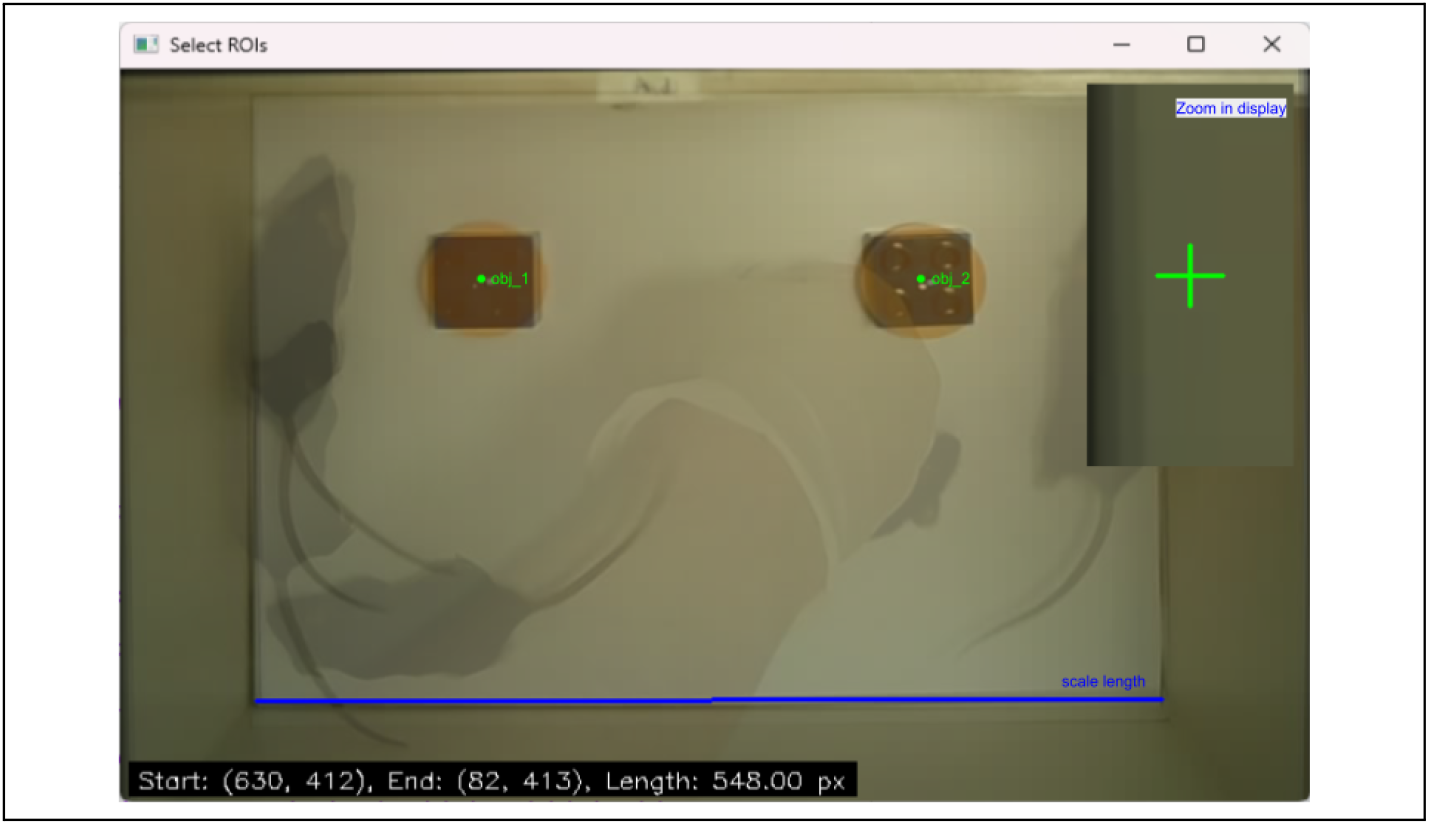
Output of the **draw_rois()** function. The background image is a composite of one frame taken from each selected video, merged to ensure that all ROI drawings align correctly across every recording.

The blue line denotes the scale length used for spatial calibration, while the saved ROI areas and points are overlaid in green.

- *Left-click to select a point*
  ○ *Press ‘S’ to save the Point*
- *Left-click and drag to draw a rectangle (Hold Ctrl while dragging to make it square)*
  ○ *Right-click and drag to move the rectangle*
  ○ *Use the scroll wheel to resize the rectangle*
  ○ *Use Ctrl + scroll wheel to rotate the rectangle*
  ○ *Press ‘S’ to save the ROI*
- *Hold Alt + left-click and drag to draw a scale line*
  ○ *Press ‘S’ to state the scale line distance (in cm)*
- *Press ‘Q’ to quit and save all ROIs*

*It also stores the scaling factor of the video (from pixels to cm) using a known distance on display. Since these ROIs will be used to analyze the mouse position, they should be drawn on the* ***final version*** *of the videos (cropped to the desired size and aligned)*.

### Basic protocol 1

#### 1. Manual Behavioral Labeling

##### Introductory paragraph

Manually scoring is a common practice in behavioral neuroscience labs, where experimenters typically watch each video and record behavioral events in real time. This process demands sustained attention for long periods of time, as any distraction often forces restarting the video. Additionally, the outcome is usually a single numeric value per behavior, making it difficult to verify or correct without re-watching the entire recording. To address these limitations, we developed the RAINSTORM Behavioral Labeler, a python-based tool that enables frame-by-frame annotation of video recordings with behavioral labels. This method significantly improves the precision, efficiency, and reproducibility of manual scoring by allowing users to make deliberate, frame-specific decisions without the constraints of real-time pressure. The labeled frames not only provide detailed insights into behavioral experiments but also serve as ground truth data for training machine learning models, paving the way for automated scoring in future analyses.

##### Necessary Resources

All the software involved in the project is Open Source and can be found on our GitHub repository. RAINSTORM is a python-based project that involves the installation of the RAINSTORM package from PyPI and running Jupyter Notebooks locally on your computer. It is recommended to use Miniconda (or Anaconda) to create a virtual environment, and running the notebooks on VS code.

This first protocol works with video files (supported formats: mp4, mkv, mov and avi), and returns csv files with the annotated behaviors on each frame. An example video file is available in the downloaded repository inside `‘docs/examples/colabeled_video’.`

##### Protocol steps with *step annotations*

###### 1.1. Open the Jupyter Notebook: 1-Behavioral_labeler.ipynb

*On VS code, you can navigate through your computer and find the Notebooks in the downloaded repository folder using the Explorer (Ctrl + Shift + E)*.

*Run the first cell to load the necessary modules, including RAINSTORM*.

###### 1.2. Run the Behavioral Labeler

*Edit the list of behaviors to be labeled, and assign a key for each*.

~~~
Behaviors= [‘obj_1’, ‘obj_2’, ‘freezing’, ‘grooming’, ‘rearing’]
keys= [‘4’,’6’,’f’,’g’,’r’] # One key for each behavior
rst.labeler()
~~~

*After running the cell, a pop up window will appear, where you need to:*

###### 1.3. Navigate to the video you want to label and select it

*If you do not have a video available on your computer, you can find a demo inside the RAINSTORM repository on* *docs/examples/colabeled_video/Example_video.mp4*.

###### 1.4. (Optional) Pick a labeled csv file

*If you have already started labeling a video, you can open the csv and continue where you left off*.

###### 1.5. Confirm the behaviors you would like to label

*As it was originally set up to label the exploration of two objects, the presets are “obj_1”, “obj_2”, “freezing”, “grooming” and “rearing”. Feel free to modify them to suit your needs*.

###### 1.6. Confirm the keyboard keys you would like to use

*One for each behavior, the presets are “4”, “6”, “f”, “g” and “r”*.

###### 1.7. Start labeling

*After a few seconds the labeler will open and the first frame will be displayed (Fig. 3). Follow the instructions on the screen to navigate through the video and label each behavior as it happens*.

**Figure 3.**
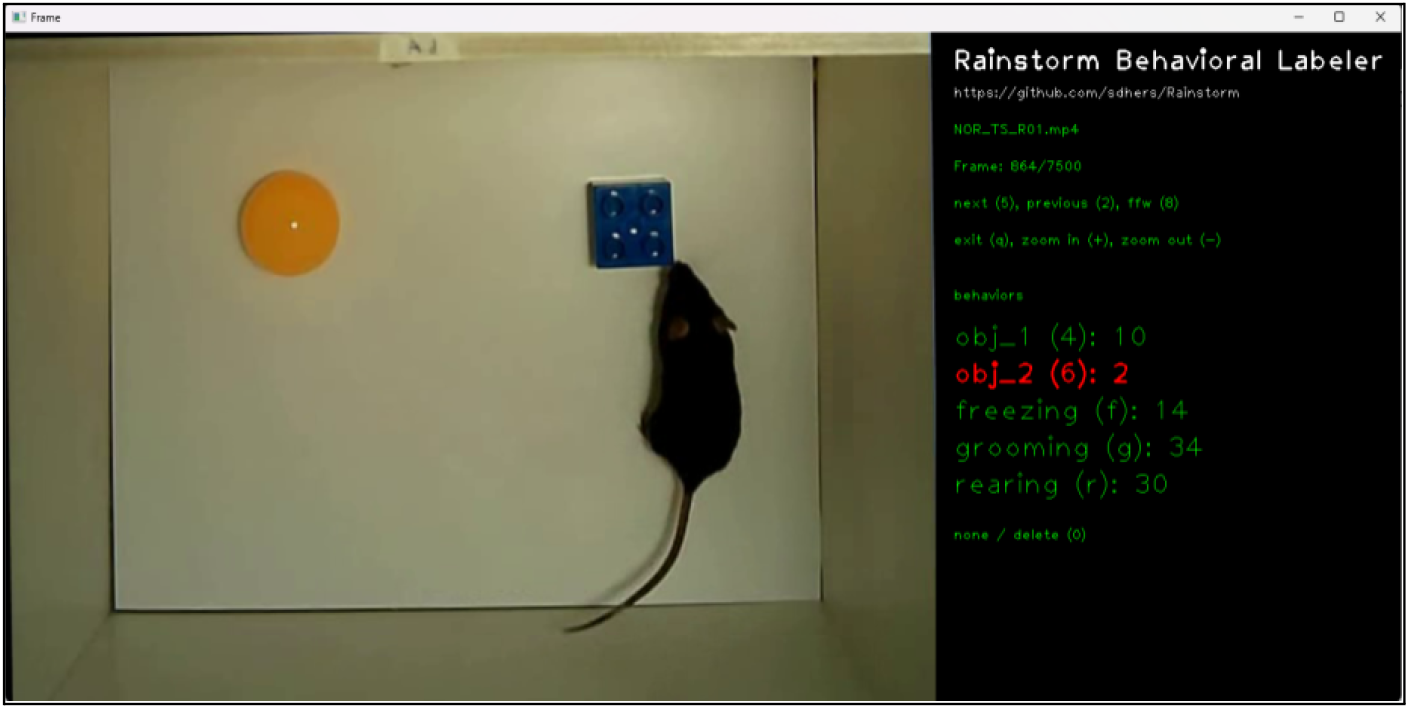
RAINSTORM Behavioral Labeler. **Left**: Current frame. **Right**: Margin with the video name, frame number, a brief summary of the navigation instructions, and a list of the selected behaviors, with their keys, and the total labeled instances for each.

Once you are done, exit and save the results. A labeled csv file will be created in the same video directory as the video.

### Basic protocol 2

#### 2. Geometric Behavioral Labeling

##### Introductory paragraph

The first approach to automate the behavioral labeling process focuses on simplifying animal movements into a series of points that track specific body parts over time. This can be achieved using pose estimation tools like DeepLabCut, which reduce the visual complexity of the animal into key points of the body. By defining exploration by the geometric relation between the points one can systematically identify and quantify it when the given geometrical rules are met (nose distance and head orientation angle towards the target are below the user defined thresholds).

##### Necessary Resources

To obtain the position of the mouse from each video, you will rely on pose estimation software. We recommend using DeepLabCut (DLC). You can find step-by-step instructions on how to install it on their webpage. The position processing also supports tracking with SLEAP (Pereira et al., 2022).

The body parts used for pose estimation will affect the behavior analysis. It is important to provide points that allow for the reconstruction of the subject’s position (Fig. 4). To use the pretrained Neural Networks in section 3.2, it is necessary to track the nose, head, neck, both ears, and body center (they do not need to have the same names, though). The custom Artificial Neural Networks will also be trained using those six points by default, but can be easily modified. Other body parts (such as shoulders, midside, hip and tail) are great for video reconstruction (see section 4.2), but are not recommended for model training.

**Figure 4.**
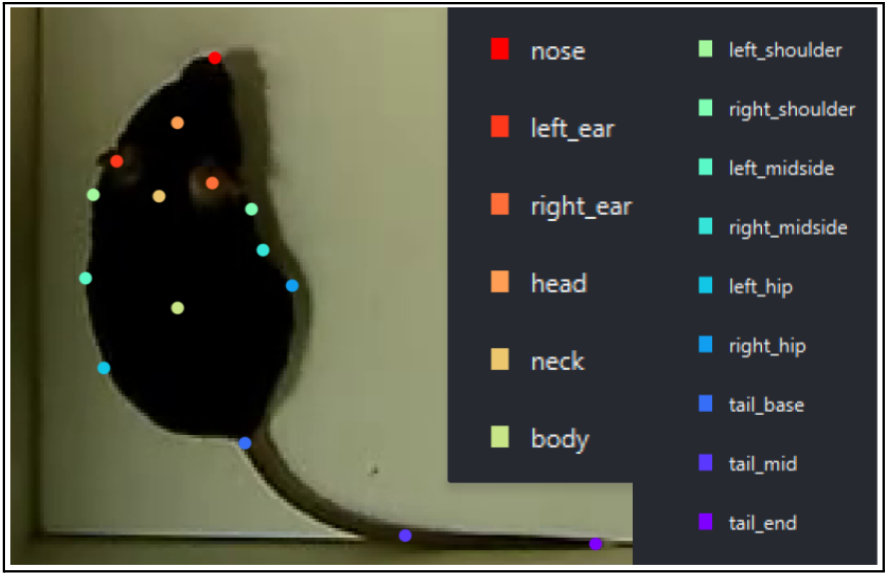
Mice body parts chosen for pose estimation. View from the DeepLabCut labeling GUI.

##### Protocol steps with *step annotations*

###### 2.1. Prepare positions

Open the Jupyter Notebook: 2a-Prepare_positions.ipynb.

Run the first cell to load the necessary modules, including RAINSTORM.

###### 2.1.1. State your project path

*You need to start by setting some variables that will adapt the processing method to our own data. The defaults are set to work with the demo data. If you are going to analyze the demo data, all you need to change is the ‘base’ variable, which should be the path to the downloaded RAINSTORM repository*.

*If you are using a Windows path with backslashes, place an ‘r’ in front of the path to avoid an error (e.g. r’C:\Users\dhers\Desktop\RAINSTORM’)*.

- *base: The path to the downloaded repository*.
- *folder_path: The path to the folder containing the pose estimation files you want to use*.
- *ROIs_path: The path to the file with the Regions of Interest (ROIs)*.

*The ROIs.json file is the one created using the ‘draw_rois’ function of the ‘0-Video_handling’ notebook*.

###### 2.1.2. Rename the files to end in ‘_position’

*The filenames that result from pose estimation analysis (e.g. DLC) end with something like* `‘{Software_used + Network + name + date + snapshot}.h5’`. *You can change the filenames on this step*.

~~~
before = ‘DLC_resnet50_shuffle2_200000.h5’
after = ‘_position.h5’
rst.rename_files(folder_path, before, after)
~~~

*(Optional) If the files belong to different trials of an experiment, they should contain the name of the trial in the filename. This will be used later to organize the files*.

###### 2.1.3. Create the params.yaml file

*The params.yaml file (params file) is a configuration file that contains all the parameters needed to run the analysis. It will be located in the experiment folder. The default parameters are preset to work with the demo files, feel free to modify them and try out different values*.

*On the params file you can modify the following parameters:*

- *path: Path to the experiment folder containing the pose estimation files*
- *filenames: Pose estimation filenames*
- *software: Software used to generate the pose estimation files (‘DLC’ or ‘SLEAP’)*
- *fps: Video frames per second*
- *bodyparts: Tracked body parts*
- *targets: Exploration targets*
- *prepare_positions: Parameters for processing positions:*
  ○ *confidence: How many std_dev away from the mean the points likelihood can be without being erased (it is similar to asking ‘how good is your tracking?’)*
  ○ *median_filter: Number of frames to use for the median filter (it must be an odd number)*
- *geometric_analysis: Parameters for geometric analysis:*
  ○ *roi_data: Loaded from ROIs.json*
    ▪ *frame_shape: Shape of the video frames ([width, height])*
    ▪ *scale: Scale of the video in px/cm*
    ▪ *areas: Defined ROIs (areas) in the frame*
    ▪ *points: Key points within the frame*
  ○ *distance: Maximum nose-target distance to consider exploration*
  ○ *orientation: Set up orientation analysis*
    ▪ *degree: Maximum head-target orientation angle to consider exploration (in degrees)*
    ▪ *front: Ending bodypart of the orientation line*
    ▪ *pivot: Starting bodypart of the orientation line*
  ○ *freezing_threshold: Movement threshold to consider freezing, computed as the mean std of all body parts over 1 second*
- *automatic_analysis: Parameters for automatic analysis:*
  ○ *model_path: Path to the model file*
  ○ *model_bodyparts: body parts used to train the model*
  ○ *rescaling: Whether to rescale the data*
  ○ *reshaping: Whether to reshape the data (set to True for RNN)*
  ○ *RNN_width: Defines the shape of the Recurrent Neural Network*
    ▪ *past: Number of past frames to include*
    ▪ *future: Number of future frames to include*
    ▪ *broad: Broaden the window by skipping some frames as it strays further from the present*
- *seize_labels: Parameters for the analysis of the experiment results:*
  ○ *groups: Experimental groups you want to compare*
  ○ *trials: If your experiment has multiple trials, list the trial names here*
  ○ *target_roles: Role/novelty of each target in the experiment*
  ○ *label_type: Label type used to measure exploration (geolabels, autolabels, etc)*

###### 2.1.4. Open an example file

*This following function will randomly choose an example file, open it and print the mean likelihood of each point in the dataframe*.

~~~
df_raw = rst.open_h5_file(params, example_path, print_data=True)
~~~

*Cell output:*

~~~
Plotting coordinates from NOR_TS_06_position.h5
Points in df: [‘body’, ‘head’, ‘nose’, …]
Body mean_likelihood: 0.98 std_dev: 0.08 tolerance: 0.82
Head mean_likelihood: 0.93 std_dev: 0.11 tolerance: 0.71
Nose mean_likelihood: 0.74 std_dev: 0.13 tolerance: 0.48
…
~~~

*Notice that, if the model is working properly, the mean likelihoods are close to 1. However, some points have lower mean likelihoods and higher standard deviations. This is because those points are harder to find (e.g. the nose tends to disappear during grooming)*.

*The Tolerance is calculated for each point as the* ***mean likelihood of the point minus the standard deviation of the likelihood multiplied by the Confidence*** *(variable set in the params file), and only the positions that are below it are erased. A higher Confidence value will then erase less points*.

###### 2.1.5. Add the position of stationary exploration targets

*The position of the exploration targets can either be tracked using the same software you use to track our animals, or you can add them here. If our pose estimation data does not have the exploration targets positions, you can add them to the DataFrame using the following ‘add_targets’ function*.

~~~
df_raw = rst.add_targets(params, df_raw, verbose=True)
~~~

*The ‘add_targets’ function will add the points from ‘roi_data’ in the params file* ***only*** *if they are also named in the ‘targets’ list on the params.yaml file*.

###### 2.1.6. Test the processing parameters on the selected video

~~~
df_smooth = rst.filter_and_smooth_df(params, df_raw)
~~~

*This cell will create a dataframe called df_smooth, that will be the result of erasing the low likelihood points (where the likelihood is below the Tolerance), interpolating, applying a median filter and gaussian smoothing to the df_raw dataframe. The function below will plot both to compare them (Fig. 5)*.

**Figure 5.**
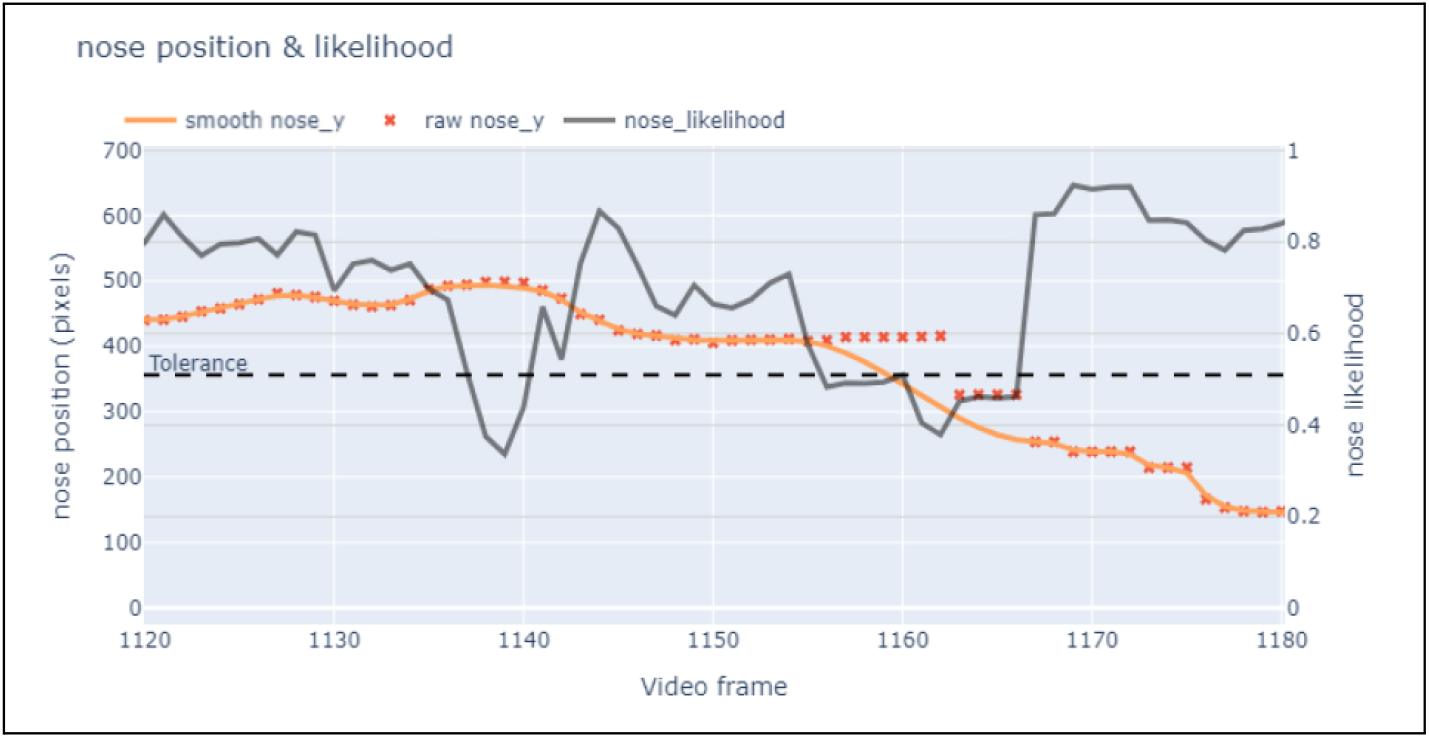
Raw vs Smoothed position of the nose: **Smooth positions** (Orange line) come from **raw positions** (Red points) after they are filtered if their corresponding **likelihood** (grey line) dropped below the **tolerance** limit (dashed line), smoothing abrupt changes resulting from brief events in which the pose estimation software is not able to recognize the recorded body part.

~~~
rst.plot_raw_vs_smoothed(params, df_raw, df_smooth, bodypart=‘nose’)
~~~

*Cell output:*

###### 2.1.7. Batch process all position files

*Once you have verified the set thresholds, you can apply all previous steps to all the files in our folder and store the results into csv files*.

~~~
rst.process_position_files(params, targetless_trials=[‘Hab’])
~~~

*Cell output:*

~~~
NOR_Hab_01_position.h5 has 30 columns. Mouse entered after 1.16 sec.
NOR_Hab_02_position.h5 has 30 columns. Mouse entered after 0.72 sec.
NOR_Hab_03_position.h5 has 30 columns. Mouse entered after 1.48 sec.
…
~~~

###### 2.1.8. Organize the resulting files into folders

*Finally, you can organize the files into subfolders corresponding to different trials of the experiment*.

~~~
rst.filter_and_move_files(params)
~~~

*Cell output:*

~~~
Files filtered and moved successfully. All .H5 files are stored away.
~~~

*The experiment folder now has subfolders according to the number of trials, each containing csv files with mice positions*.

#### 2.2. Geometric analysis

You can move on to the next notebook, 2b-Geometric_analysis.ipynb:

##### 2.2.1. State your project path & thresholds

*You need to define the path to the same folder used in ‘2-Prepare_positions’, and the path to the parameters file (which contains the thresholds for the geometric analysis)*.

*This notebook will use variables from the ‘params’ file that will adapt the processing method to our own data. The default values are set to work with the demo files*.

- *distance: Maximum nose-target distance to consider exploration*.
- *angle: Maximum nose-head-target orientation angle to consider exploration*.
- *freezing_threshold: Movement under which it is considered freezing, calculated as the mean standard deviation for all body parts over one second*.

*Also, ROI data will be used to plot and calculate time spent on each of the drawn areas*.

##### 2.2.2. Plot the trajectory of a position file

*Choose an example file, plot the nose trajectory and visualize the thresholds (Fig. 6). The function will label exploration events when the nose is both colored (heading towards the target) and inside the dashed line (close to the target)*.

**Figure 6.**
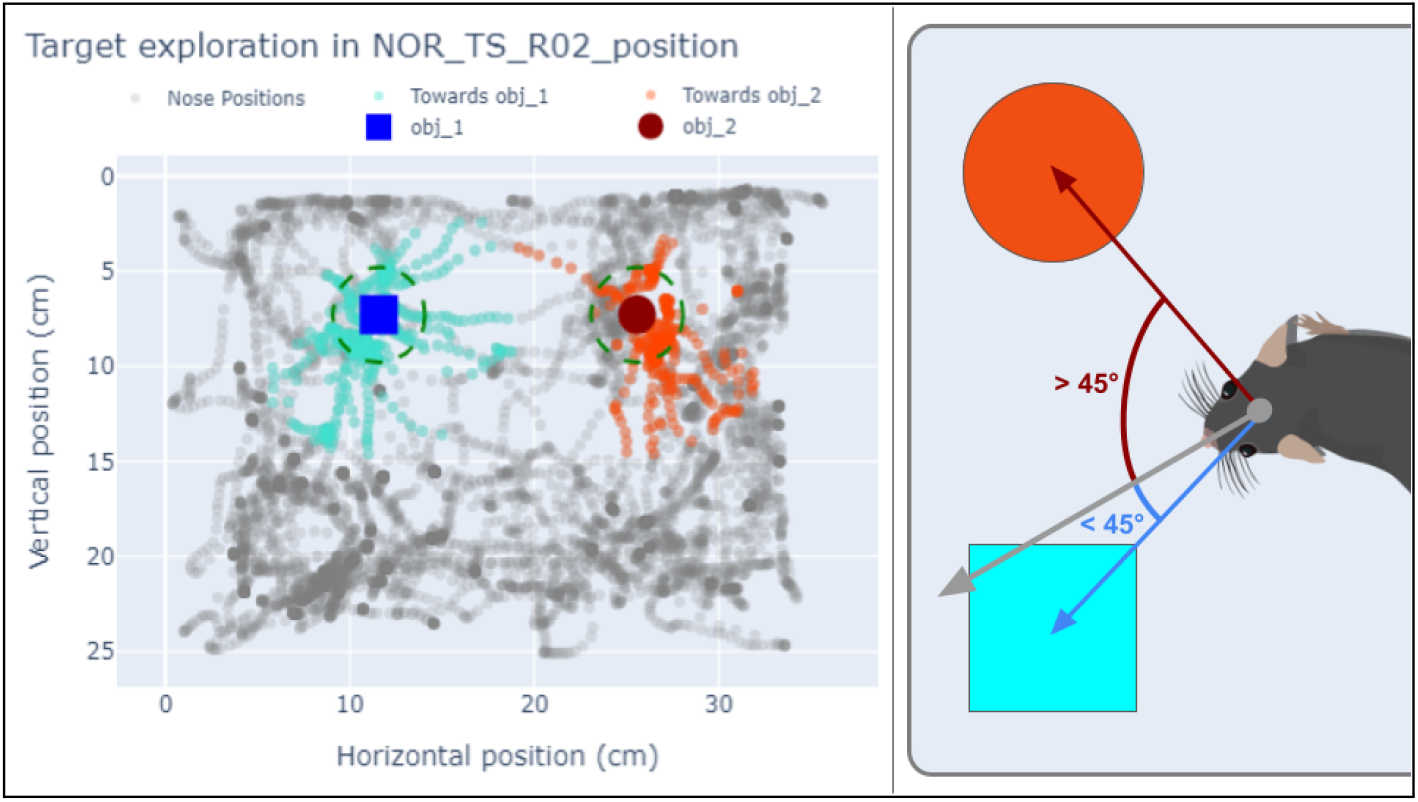
**Left**: Trajectory of the nose during exploration. Each point is a position occupied by the nose at some time during the session. The nose position is colored when it is oriented towards one of the targets (blue square or red circle, default nose-head-target angle is 45°), and a dashed green line shows the distance nose-target up to which the method will consider the animal is exploring (default is 2.5 cm). **Right**: Illustration of nose-head-target angle computation. The gray arrow depicts the head-to-nose vector, and the blue arrow depicts the head-to-target vector toward the blue square. The angle between these two vectors is less than 45°, classifying the mouse as oriented toward that object (mouse drawing adapted from Claudi, 2020).

~~~
Rst.plot_position(params, example_path)
~~~

*Cell output:*

##### 2.2.3. Use the positions to measure time spent in each ROI

Measure the time spent on each of the areas drawn using the draw_rois() tool on section 0.7. `roi_activity = rst.detect_roi_activity(params, example_path)`

~~~
rst.plot_heatmap(params, example_path)
~~~

##### 2.2.4. Measure freezing

Compute and plot the mouse’s overall movement as the standard deviation of frame-to-frame displacements over a one-second rolling window. By applying a movement threshold (default = 0.01 cm/s, adjustable in the ‘params’ file), periods below that threshold are classified as freezing, allowing us to quantify total freezing time.

~~~
rst.plot_freezing(params, example_path)
~~~

##### 2.2.5. Analyze all files in the experiment folder

*The following function will use the previously set thresholds to create two files for each position file: A ‘movement.csv’ file containing distance traveled, freezing and darting; and a ‘geolabels.csv’ file with the exploration of each target, frame by frame*.

~~~
rst.create_movement_and_geolabels(params, roi_bodypart = ‘body’)
~~~

*Cell output:*

~~~
Saved geolabels to NOR_TS_01_geolabels.csv
Saved movement to NOR_TS_01_movement.csv
Saved geolabels to NOR_TS_02_geolabels.csv
…
~~~

This is the end of the Geometric Analysis protocol.

### Basic protocol 3

#### 3. Automated Behavioral Labeling

##### Introductory paragraph

This automated labeling protocol is a major step forward, using advanced machine learning models to simplify behavioral analysis. While manual labeling and algorithmic filtering based on positional data offer valuable insights, this new approach blends expert-defined criteria with precise pose estimation parameters to train Artificial Neural Networks (ANN) that accurately identify and label behavior. We focus on labeling exploratory behavior because it is directly related to memory performance in our studies, though the method is versatile and can be applied to all other observable behaviors present in the recordings.

In addition, the protocol incorporates Recurrent Neural Networks (RNN, or ‘wide’ models) to analyze sequences of frames, capturing temporal context (past and future relative to the frame being analyzed) to better understand behavior. By automating this process, the protocol significantly reduces manual workload while maintaining high accuracy, making it ideal for large-scale studies in neuroscience, ethology, and other fields. Its adaptability and robustness ensure seamless integration into diverse experimental protocols, paving the way for more comprehensive and efficient behavioral research.

##### Necessary Resources

The creation and training of Artificial Neural Networks require both positional data and corresponding labeled behaviors for the same set of videos. To facilitate this, we provide a dataset containing positional information for over a hundred video fragments, each labeled by five experimenters with varying levels of expertise. For those wishing to create their own training dataset, behavioral labels can be obtained manually by following the Manual Behavioral Labeling Protocol, while pose estimation can be performed using DLC. The resulting position data can then be processed with the Geometric Behavioral Labeling Protocol for smoothing before training the models.

While the neural networks can be trained using a CPU, the process is significantly faster with a GPU. It is recommended to use a GPU setup with CUDA (Compute Unified Device Architecture) compatible with TensorFlow 2.10 (ensuring compatibility with DLC v2.0 as well).

##### Protocol steps with *step annotations*

###### 3.1. Create Models

*Open the Jupyter Notebook: 3a-Create_models.ipynb*

###### 3.1.1. State your project path

*Define the path to the folder where the trained models will be stored*.

###### 3.1.2. Create the modeling.yaml file

*The modeling.yaml file is a configuration file that contains all the parameters needed to create and train the models. It will be located in the models folder*.

~~~
modeling = rst.create_modeling(models_folder)
~~~

*It contains the following parameters:*

- *path: Path to the models folder*
- *colabels: The colabels file is used to store and organize positions and labels for model training*
  ○ *colabels_path: Path to the colabels folder*
  ○ *labelers: List of labelers on the colabels file (as found in the columns)*
  ○ *target: Name of the target on the colabels file*
- *focus_distance: Window of frames to consider around an exploration event*
- *bodyparts: List of body parts used to train the model*
- *split: Parameters for splitting the data into training, validation, and testing sets*
  ○ *validation: Percentage of the data to use for validation*
  ○ *test: Percentage of the data to use for testing*
- *RNN: Set up the Recurrent Neural Network*
  ○ *width: Defines the shape of the wide model*
    ▪ *past: Number of past frames to include*
    ▪ *future: Number of future frames to include*
    ▪ *broad: Broaden the window by skipping some frames as it strays further from the present*
  ○ *units: Number of neurons on each layer*
  ○ *batch_size: Number of training samples the model processes before updating its weights*
  ○ *epochs: Each epoch is a complete pass through the entire training dataset*
  ○ *lr: Learning rate*

###### 3.1.3. Prepare the training data

*First, load the dataset from the colabels file and create one ‘labels’ column out of all the labelers*.

~~~
dataset = rst.prepare_data(modeling)
~~~

*Next (optional, but recommended), erase the rows of the dataset that are too far away from exploration events*.

~~~
dataset = rst.focus(modeling, dataset)
~~~

*Finally, split the dataset into training, testing and validation subsets*.

~~~
model_dict = rst.split_tr_ts_val(modeling, dataset)
~~~

*The model_dict contains the training, testing and validation datasets for both simple and wide (which include a temporal window of frames) models*.

###### 3.1.4. Train the first ‘simple’ model

*With the training data ready, use TensorFlow to design your very first ANN*.

- *It will look at the positions of one frame at a time, and try to decide if the mouse is exploring. If the decision is correct the architecture will be reinforced, else it will be corrected according to the learning rate*.
- *It will train for some epochs (cycles through the whole dataset) and plot how the accuracy and loss (error between the predictions and the target values) evolve (Fig. 7)*.
- *You will be validating the training using the validation split, which contains frames that were not used for training*.

**Figure 7.**
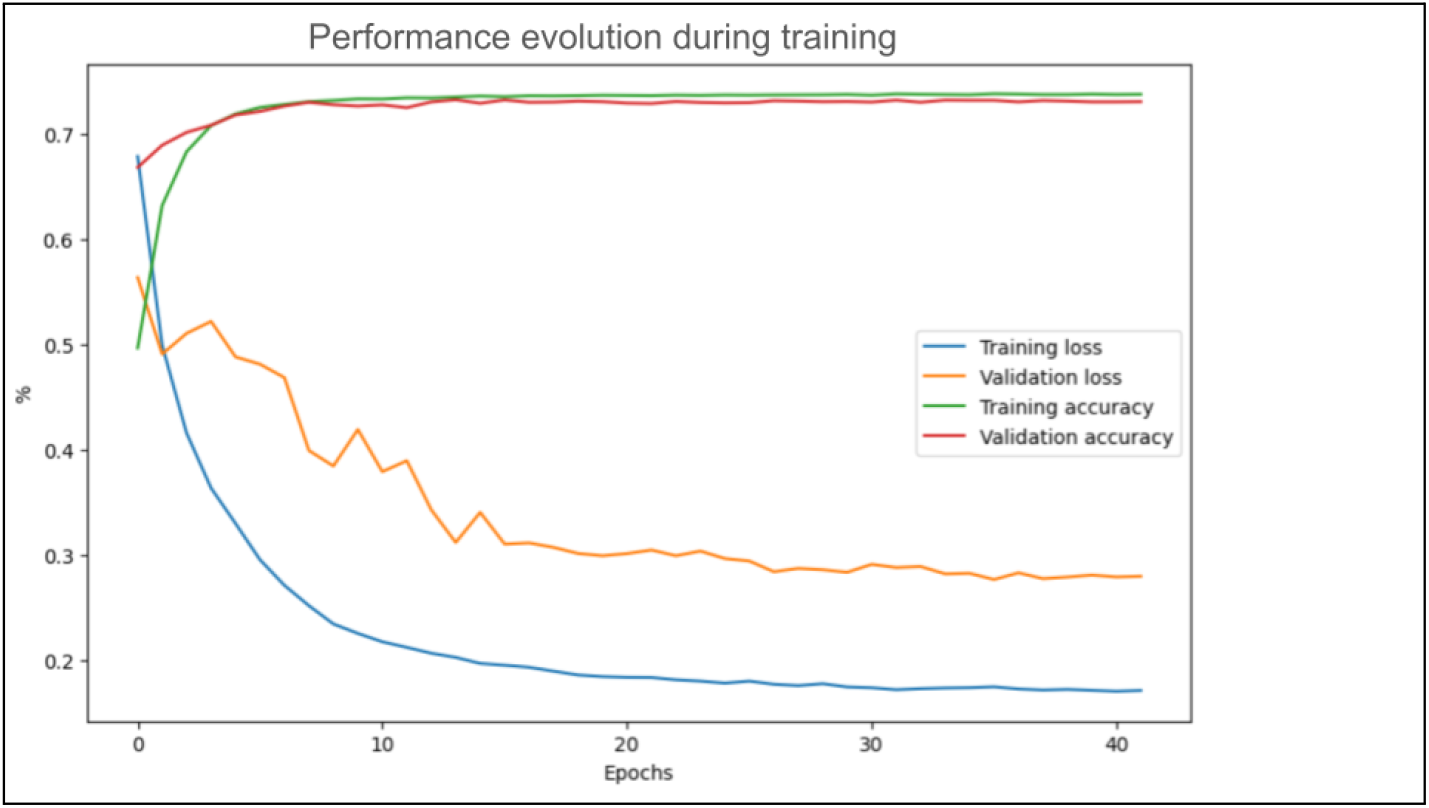
Training of an ANN. Both accuracy and loss are calculated on the training dataset (green and blue lines) to ensure the model is learning correctly. They are also calculated on the validation dataset (red and orange lines), which contains positions not used for training, to prevent overfitting.

~~~
model_simple = tf.keras.Sequential([
Input(shape=(X_tr.shape[1],)), # Input layer
Dense(32, activation=‘relu’),
…
Dense(8, activation=‘relu’),
Dense(1, activation=‘sigmoid’)])
model_simple.compile(optimizer=tf.keras.optimizers.Adam(learning_rate = 0.0001), loss=‘binary_crossentropy’, metrics=[‘accuracy’])
history_simple = model_simple.fit(X_tr, y_tr, epochs=10, batch_size=64, validation_data=(X_val, y_val))
rst.plot_history(history_simple, “Simple”)
~~~

*Cell output:*

###### 3.1.5. Train a Recursive Neural Network

*Now that you have a simple model trained, you can start building more complex models with the help of some functions. To make our artificial networks as real as possible, you can let them see a sequence of frames to decide if the mouse is exploring*.

- *RAINSTORM’s build_RNN function will use Bidirectional Long Short-Term Memory (LSTM) layers that allow the model to take into account the sequence of frames*.
- *With the ‘units’ parameter you can say how many and how big our hidden layers will be*. `model_wide = rst.build_RNN(modeling, model_dict)`
- *The train_RNN function includes ‘early stopping’ and ‘learning rate scheduler’ mechanisms that will prevent the model from overfitting*.

~~~
history_wide = rst.train_RNN(modeling, model_dict, model_wide)
~~~

*Cell output:*

~~~
Epoch 1/60
423/423 [==============================] - 13s 13ms/step - loss: 0.7437 -
accuracy: 0.5096 - val_loss: 0.4357 - val_accuracy: 0.7552 - lr: 6.0000e-05 Epoch 2/60]
423/423 [==============================] - 4s 9ms/step - loss: 0.5570 -
accuracy: 0.6338 - val_loss: 0.3426 - val_accuracy: 0.7898 - lr: 7.3333e-05 Epoch 3/60
…
Epoch 32/60
418/423 [============================>.] - ETA: 0s - loss: 0.1769 - accuracy:
0.7990
Restoring model weights from the end of the best epoch: 26.
423/423 [==============================] - 4s 10ms/step - loss: 0.1770 -
accuracy: 0.7984 - val_loss: 0.1693 - val_accuracy: 0.8271 - lr: 1.4358e-05 Epoch 32: early stopping
~~~

###### 3.1.6. Compare the ‘simple’ and ‘wide’

*Finally, it is time to compare all the trained models. You have trained using the training dataset, and validated using the validation dataset. Now, introducing no plot twist, you will test each model using the testing dataset*.

`precision, recall, f1, mse, mae, r2 = rst.evaluate(y_pred, y_ts)`

*Cell output:*

`Precision = 0.9302, Recall = 0.9271, F1 Score = 0.9281, MSE = 0.0392, MAE = 0.0983, R-squared = 0.7681 -> simple`

`Precision = 0.9325, Recall = 0.9323, F1 Score = 0.9324, MSE = 0.0351, MAE = 0.0755, R-squared = 0.7921 -> wide`

*Metrics:*

- *Precision - Of all the positive predictions made, how many were actually correct*.
- *Recall (or Sensitivity) - Of all the actual positive cases, how many were correctly predicted*.
- *F1 Score - The harmonic mean of Precision and Recall, balancing between correctly identifying positives and avoiding false positives*.
- *Mean Squared Error - Measures the average squared difference between predicted and actual values*.
- *Mean Absolute Error - Measures the average absolute difference between predicted and actual values*.
- *R-squared - Indicates how well the model explains the variance in the data*.

*Our trained models are stored safely in our repository, with today’s date. You can now compare each model with the performance of the human labelers*.

###### Evaluate models

*One may think that the evaluation on the testing set is enough, and in many cases it is. However, for the purpose of finding a model that resembles the labeling of an expert, It is better to compare the performance of the model against all the manually labeled data*.

###### 3.1.7. Create a good reference to compare against

*Since you want to compare the models and the labelers, you need to create an appropriate reference. This reference could be the mean of all the labelers, but then the labelers would have an unfair advantage.To avoid this, the following function will simultaneously create a chimera labeler and a leave-one-out-mean:*

- *The chimera is created by randomly selecting a labeler on each row of the data*.
- *The leave-one-out-mean is created by averaging the remaining values*.

*This way, you can compare the chimera to the leave-one-out mean knowing that they are independent*.

~~~
chimera, loo_mean = rst.create_chimera_and_loo_mean(manual_labels, seed=42)
~~~

###### 3.1.8. Load the models

*List the models you want to evaluate:*

~~~
model_paths = {
‘example_simple’: os.path.join(models_folder, ‘trained_models/example_simple.keras’), ‘example_wide’: os.path.join(models_folder, ‘trained_models/example_wide.keras’),
…}
~~~

*Use each model to label exploration on all the available data:*

~~~
models_dict = rst.build_and_run_models(modeling, model_paths, evaluation_dict[‘position’])
~~~

###### 3.1.9. Evaluate the performance metrics

*You can compare the predicted labels from each model against the leave-one-out mean as well as the chimera and each of the human labelers separately (Table 2). Remember that the chimera is independent from the leave-one-out mean, but the labelers are not*.

**Table 2.** Metrics of performance for each method. Each model or human labeler was evaluated against the leave-one-out mean, which is independent from the chimera labeler. Darker color stands for better performance.

| Model | Precision | Recall | F1 Score | MSE | MAE | R-squared |
| --- | --- | --- | --- | --- | --- | --- |
| Simple | 0.9826 | 0.9801 | 0.9809 | 0.0102 | 0.0262 | 0.8072 |
| Wide | 0.9865 | 0.9854 | 0.9858 | 0.0070 | 0.0204 | 0.8671 |
| Chimera | 0.9795 | 0.9781 | 0.9786 | 0.0177 | 0.0239 | 0.6645 |
| Geometric | 0.9732 | 0.9746 | 0.9726 | 0.0231 | 0.0293 | 0.5628 |
| Labeler_A | 0.9846 | 0.9800 | 0.9812 | 0.0126 | 0.0188 | 0.7613 |
| Labeler_B | 0.9802 | 0.9801 | 0.9783 | 0.0185 | 0.0246 | 0.6506 |
| Labeler_C | 0.9786 | 0.9780 | 0.9750 | 0.0191 | 0.0252 | 0.5633 |
| Labeler_D | 0.9857 | 0.9845 | 0.9849 | 0.0125 | 0.0186 | 0.7634 |
| Labeler_E | 0.9900 | 0.9902 | 0.9900 | 0.0082 | 0.0143 | 0.8454 |

*Regardless of the metrics obtained by each, one should aim for the models to be similar to the mean of the labelers. You can test that by running a Principal Components Analysis (PCA) on the labels and plotting them to measure the distance between each method and the mean (Fig. 8)*.

**Figure 8.**
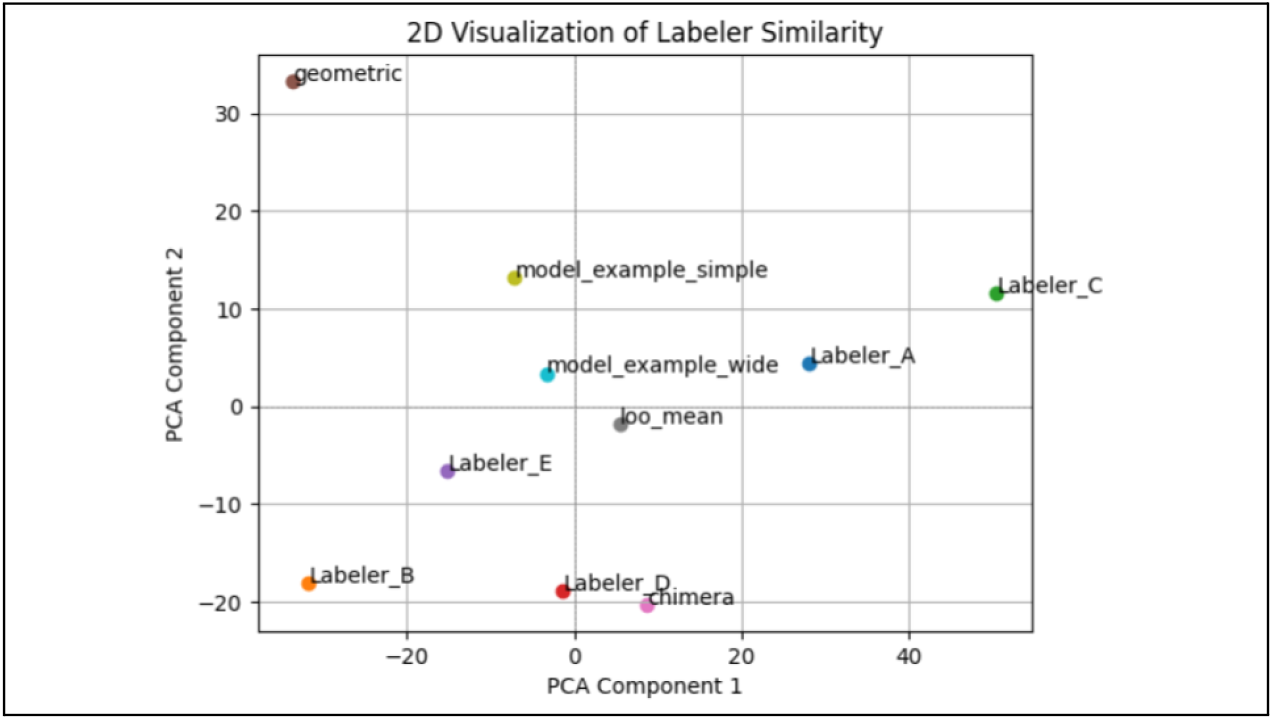
PCA of labeling-method distances from the mean. Vectorized labels are projected onto new axes, called principal components, that maximize the variance of the data. The models come closer to the mean as they increase in complexity, while human labelers (A to E) are scattered along both axes.

~~~
rst.plot_PCA(all_labelers)
~~~

*Cell output:*

*Also, you can see both the models and the labelers performance on an example video (Fig. 9):*

**Figure 9.**
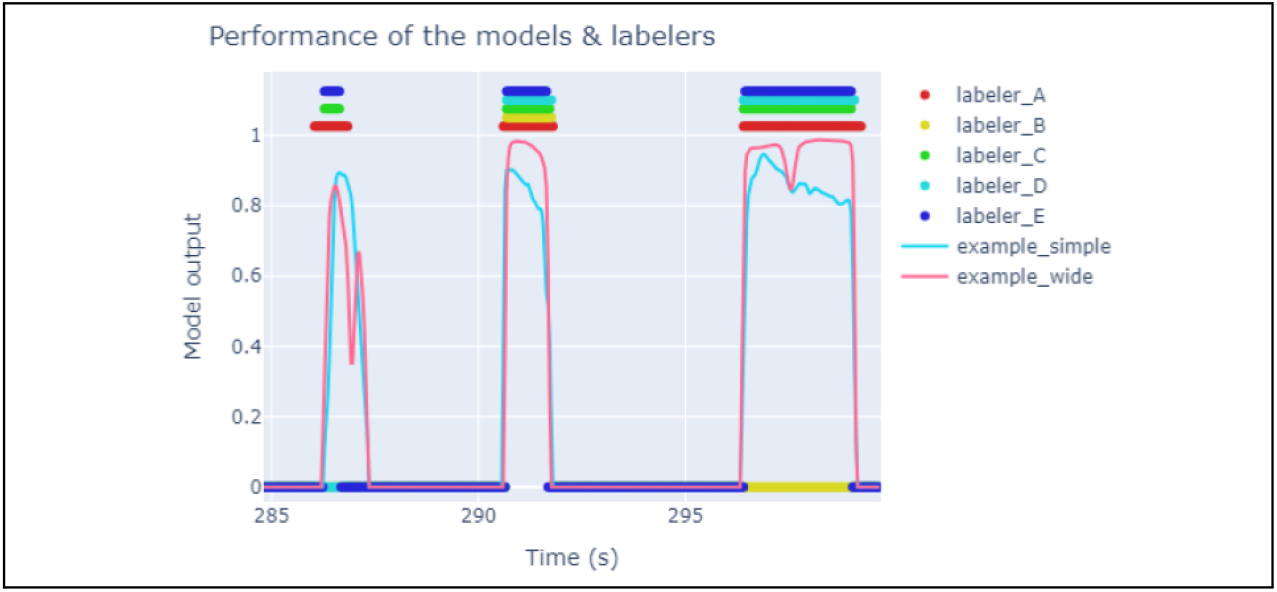
Comparison of each ANN trained model output on an example video timeline. **Wide** (red line) and **Simple** (green line) models are plotted as thin lines, labelers criteria are plotted as thick lines.

~~~
rst.plot_performance_on_video(example_path, models, labelers_example,
plot_obj=“obj_2”)
~~~

*Cell output:*

*At this point, you can select your favorite model. Next, you can move on to the following Notebook and use the chosen model to create behavioral labels for the position files*.

###### 3.2. Automatic analysis

Open the jupyter notebook: 3b-Automatic_analysis.ipynb

###### 3.2.1. State your project path, and set up the model structure

*To make sure the data will be shaped as the one used for training the models, set on the params.yaml file the same Width (past, future, broad) you had on the modeling.yaml file*.

Analyze all files in the experiment folder.

*Based on what you evaluated previously, you can choose the model of preference to analyze the position files. Set the path to the chosen model on the params file*.

*Once you run this cell, the model will be used on each of the position files on the experiment folder and you will obtain an ‘autolabels.csv’ file for each file, with columns corresponding to each of the targets:*

~~~
rst.create_autolabels(params)
~~~

Cell output:

~~~
467/467 [==============================] - 1s 3ms/step
Saved autolabels to NOR_TR_C1_D_autolabels.csv
464/464 [==============================] - 1s 3ms/step
Saved autolabels to NOR_TR_C2_A_autolabels.csv
…
~~~

**Compare labels:**

*By following the pipeline as it is presented here, upon reaching this final part there will be Manual, Geometric and Automatic labels for target exploration. Those files will be the ones needed to compare and plot. For those who have followed the analysis steps with the example files provided on the repository, we provide the Manual labels for those example videos (obtained using the RAINSTORM Behavioral Labeler). Original videos are provided upon contact in order to label them and compare experimenter criteria with the machine*.

###### 3.2.3. Open an example position file and plot its labels

~~~
positions, manual_labels, geolabels, autolabels = rst.compare_labels(folder_path)
~~~

*A great way of visualizing the distance and angle of approach to a target is to use a polar graph (Fig. 10). The distance is represented in the radius of the circle, and the circumference represents the angle of the vector from the head to the nose*.

**Figure 10.**
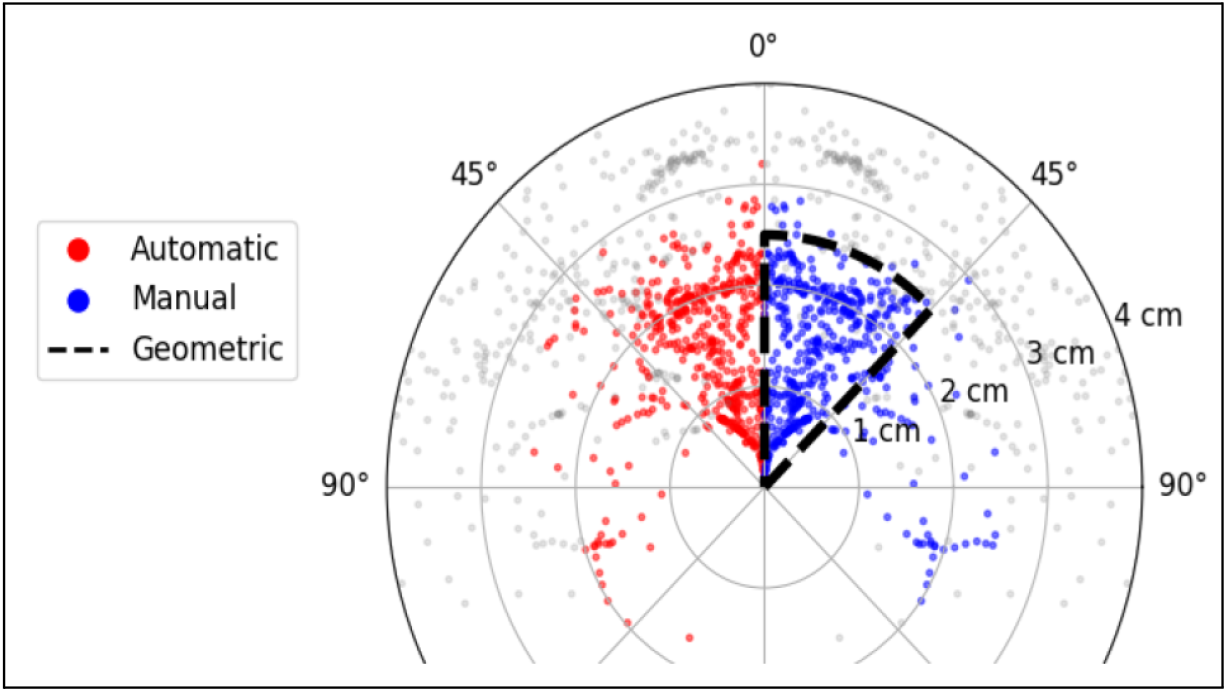
Comparison of each method on a Polar Graph. Since the graph is symmetrical, the left side will be used to color the **automatic labels** (red), and the right side to color the **manual labels** (blue). The graph shows the **geometric labels** as all the points that fit inside the dashed lines.

*Since the graph is symmetrical, the function will use the left side to color the automatic labels in red, and the right side to color the manual labels in blue. The graph will also show the geometric labels as all the points that fit inside the dashed line*.

~~~
rst.polar_graph(params, positions, autolabels, manual_labels)
~~~

*Cell output:*

###### 3.2.4. Evaluate accuracy of predictions

*Finally, you can evaluate the accuracy of the predictions by comparing the geometric and automatic labels to the manual labels*.

~~~
rst.accuracy_scores(all_manual_labels, all_geolabels, …
~~~

*Cell output:*

Mice explored **6.96%** of the time. The **geometric method** measured **6.12%** of the time as exploration. It got **25.96%** of false negatives and **13.82%** of false positives. The **automatic method** measured **7.11%** of the time as exploration. It got **12.85%** of false negatives and **8.55%** of false positives.

*Both geometric and automated labeling methods exhibit errors when compared to manual annotations. However, manual annotations themselves are subject to the biases and variability of a single observer. By aggregating the labels of several independent human evaluators to form a consensus reference, a more robust ground truth can be obtained and thus reduce the apparent error of geometric and automated methods*.

### Basic protocol 4

#### Plotting Behavioral Labels

##### Introductory paragraph

One of the most critical aspects of behavioral analysis is the ability to visually compare and evaluate the obtained results in an intuitive manner. Before conducting statistical analyses, it is essential to assess whether the recorded data reveals any unexpected outcomes, such as immobile mice or abnormal behaviors. In this final section you will find tools to visualize and explore the results derived from the Manual, Geometric, and Automated Behavioral Labeling methods. These visualizations enable users to compare exploration dynamics across experimental groups using line plots, histograms, and boxplots for metrics like total time and discrimination index. Additional plots include—but are not limited to—total distance traveled, time spent freezing, histograms of freezing event duration, and darting behaviors.

##### Necessary Resources

If one follows the pipeline as it is described before, the requirements for this last protocol are stored inside the experiment folder. To plot the experiment results, you need to have analyzed the positions and labeled the behavior of each mouse. You are ready to assign an experimental group to each animal and plot the experiment results.

##### Protocol steps with *step annotations*

###### Seize Labels

Open the jupyter notebook: 4-Seize_labels.ipynb

###### 4.1 State your project path and experiment details

*Start by setting some variables that will organize the plots according to the experimental groups. The defaults are set to work with the demo files*.

- *base: The path to the downloaded repository*.

*Go to the params.yaml file and specify the following parameters in the ‘seize_labels’ section:*

- *groups: Name the groups on your experiment*.
- *trials: Name the trials on your experiment*.
- *target_roles: Role/novelty of each target in the experiment*.
- *label_type: State which labels you want to use*.

###### 4.2. Visualize the behavioral labels on a video

*The following function combines the position data, the behavioral labels, and the original video recording to create a video with the labels overlaid (Video 1)*.

~~~
rst.create_video(params, example_path, video_path)
~~~

**Video 1.**
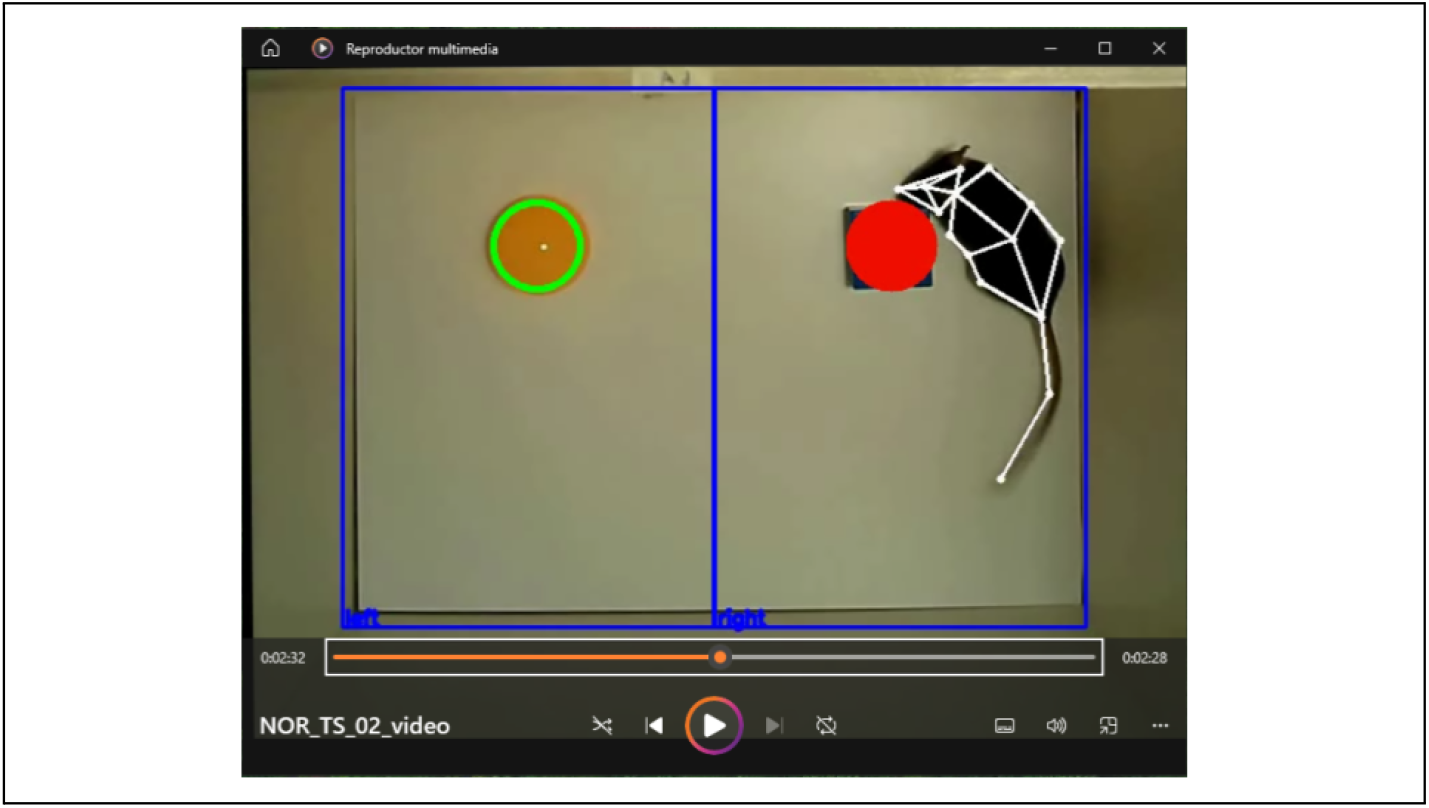
Overlay of the original video recording, the mouse pose estimation tracking data, the selected ROIs and the target’s positions. Targets change color from green to red according to the chosen labels to show when the method labeled exploration.

*The video can be built both using the original video file or not*… *try removing the ‘video_path’ argument from the function call to see the difference*.

###### 4.3. Plot individual mouse exploration

*Exploration is a dynamic behavior, and experiment results can change a lot depending on the timeframe chosen for analysis. A first step to understanding the exploration dynamics is to plot the evolution of the cumulative exploration time (Fig. 11)*.

**Figure 11.**
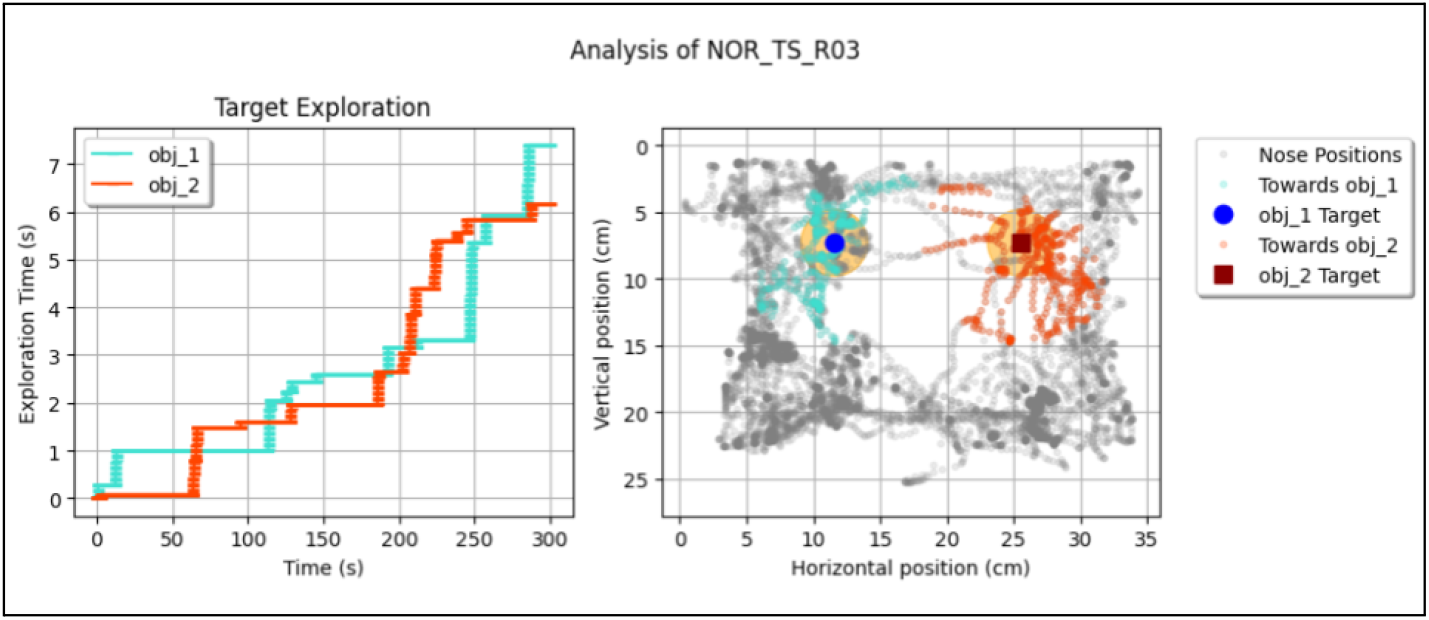
Individual exploration plot. **Left**: Cumulative exploration time for each target. **Right**: Trajectory of the subject’s nose during exploration. As in fig. 6A, the nose is colored when it is oriented towards one of the targets (default angle is 45°), and the orange area shows the nose-target distance up to which the geometric model considers exploration (default is 2.5 cm).

~~~
rst.plot_mouse_exploration(params, example_path)
~~~

*Cell output:*

*This plot shows the behavior of a single subject, and does not take into account the role of each exploratory target (i.e. which target is familiar and which is novel). To build more elaborate plots, you need to inform the groups and target roles for each subject*.

###### 4.4. Create and complete the ‘reference’ file

*This function will use the groups, trials and target roles you stated on the params file to create a ‘reference.csv’ file that will be used to organize the files and plot the experiment results*.

~~~
reference_path = rst.create_reference_file(params)
~~~

*Cell output:*

~~~
‘C:\Users\dhers\RAINSTORM\docs\examples\NOR_example_copy\reference.csv’ created successfully with the list of video files.
~~~

*Go to the experiment folder and complete the ‘reference.csv’ file. If you are using the NOR example folder to try out RAINSTORM, you will find a copy of the ‘reference.csv’ file* ***already completed*** *in the experiment folder*.

*With the ‘reference.csv’ file complete, you can proceed to the next step where it will be used to create the ‘summary’ folder*.

###### 4.5. Create the ‘summary’ folder

*The groups and targets on the ‘reference’ file will be used to organize the files into subfolders, and the target exploration columns will be renamed according to their role (e.g. Novel and Known)*.

~~~
summary_path = rst.create_summary(params)
~~~

*Cell output:*

~~~
Renamed and saved NOR_TS_C1_A_summary.csv
…
~~~

###### 4.6. With the “summary” files ready, run the analysis and plot the results

~~~
rst.plot_multiple_analyses(params, ‘TS’,[
rst.lineplot_exploration_cumulative_time,
rst.boxplot_exploration_cumulative_time, rst.plot_DI,
rst.lineplot_cumulative_distance,
rst.lineplot_freezing_cumulative_time
])
~~~

*Cell output:*

*These modular graphs (Fig. 12) are intended to show the dynamic behavior of mice, providing a deeper insight into the final result of the experiment (learning and memory in the example provided here)*.

**Figure 12.**
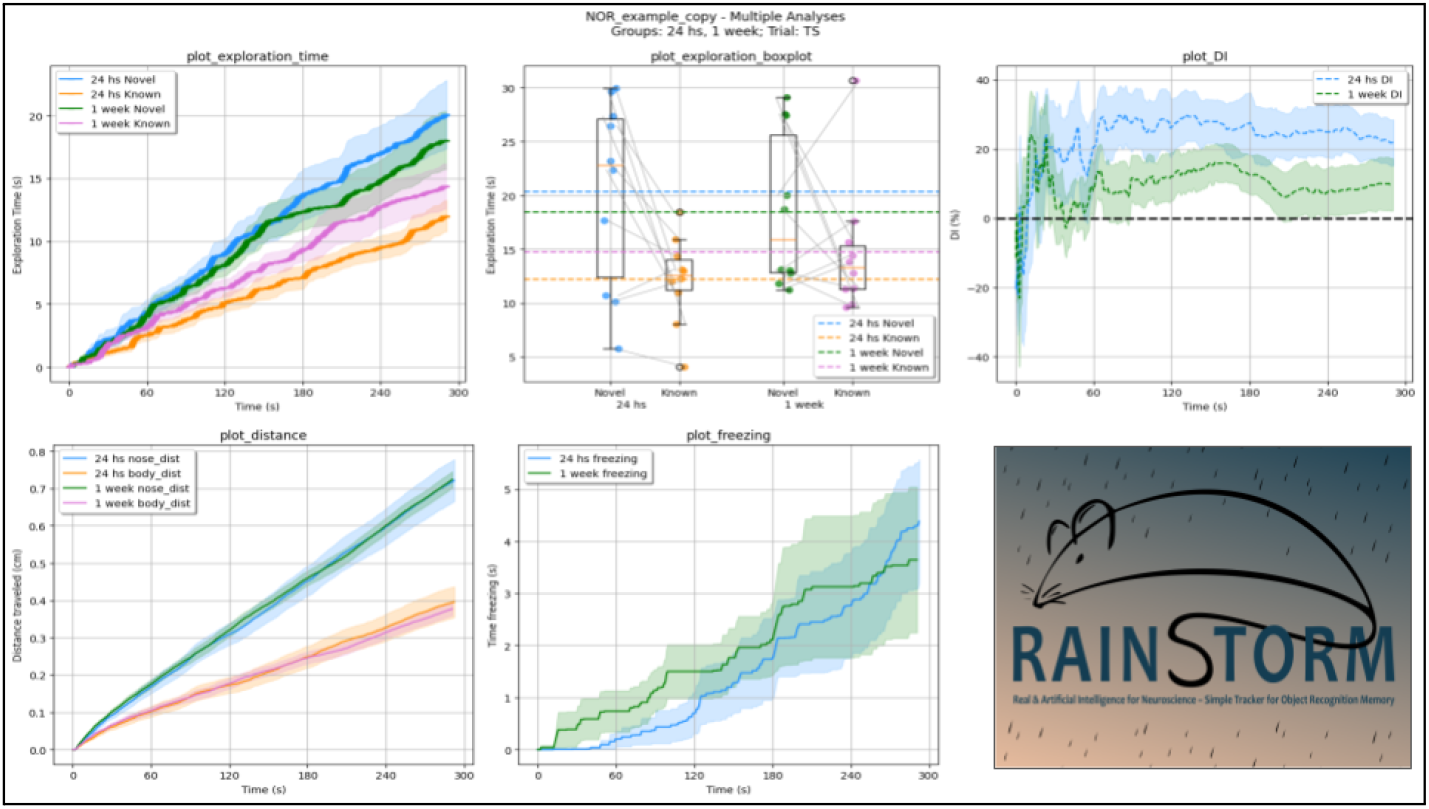
Collection of Behavioral plots describing a Novel Object Recognition testing session. From left to right, top to bottom: Cumulative exploration time (s) for each tested target at two different intervals after sample/training phase (24 h and 1 wk); Exploration time boxplots (s) of the whole session; Instant discrimination index (DI, %) throughout the session; Cumulative distance (cm) traveled; and Cumulative time spent freezing (s) during the session.

### Guidelines for understanding results

Results from processing trajectories with pose estimation software are greatly influenced by the accuracy and consistency of the raw position data. The quality of the pose estimation data depends heavily on both the precision of the model and the size of the dataset used during training. In cases where pose estimation is poor, trajectories can exhibit glitches such as sudden jumps or missing body parts. To mitigate these issues, it is advisable to process data points with lower **confidence** values (see Section 2.1.6) and ensure that any interpolation performed excludes positions with low likelihood, as including them may adversely affect subsequent analysis steps.

When evaluating RAINSTORM-trained models, it is important to interpret the results with careful attention to key statistical indicators and quality control measures. Expected statistical values—such as **precision, recall**, and **F1 score**—should ideally be higher than those obtained from individual labelers, and in the best-case scenario, even surpass the performance of the chimera labels. The close distance between the mean manual and automated labels in the **PCA analysis** strongly suggests that the training process succeeded. All indicators aside, the most important validation is given by the visual inspection of the tracked positions and behavioral classifications (see figures 9 and 10). Flags indicating unreliable results may include an unusually high number of frames classified as “exploring”, and could be indicating an imbalance in the representation of exploratory vs non exploratory frames in the training dataset which can be adjusted by using the focus function on section 3.1.3.

Reasonable conclusions drawn from RAINSTORM analyses include quantification of object exploration patterns, comparison of exploration behavior across experimental groups, and identification of trends over time. However, researchers should take care not to overinterpret subtle differences, especially when the results are not statistically significant. Line plots including the mean response for a group of animals also include the standard error to help visualize possible differences between groups, but do not assure statistical differences. If results are inconsistent with prior behavioral expectations (e.g., a complete lack of object preference in a well-characterized experimental setup), it may be necessary to revisit preprocessing steps, such as mice and targets pose estimation, or model training. Even when the obtained results align with the expectations, it is important to consider that the automatic analysis does not fully replace the experimenter’s visualization of the behavior, as RAINSTORM is not able to detect subtle irregular or atypical behaviors that could go unnoticed.

### Commentary

#### Background Information

Accurately quantifying exploratory behavior in rodents is essential for studying cognitive processes like object recognition memory. Traditional manual scoring is time-consuming, prone to bias, and difficult to standardize, while commercial alternatives — although user-friendly — often impose high costs and restrict customization. The **Geometric behavioral labeling** protocol offers a customizable approach similar to commercial tools, allowing users to set distance and angle thresholds. However, even with fine-tuning, its results deviate more from expert criteria than the Automatic protocol (See figures 8 and 10). Compared to manual scoring and threshold-based tracking, **RAINSTORM’s RNN models** closely align with expert criteria, achieving high precision and scalability.

RNNs are particularly well-suited for this task because they capture the temporal dynamics and long-range correlations inherent in behavioral data. Unlike conventional methods that require hand-engineered features, these models learn directly from raw data, enabling them to accurately model complex, time-sequence dependent behaviors. Recent studies (Mienye et al., 2024) have demonstrated that such architectures not only predict future actions with high accuracy but also reveal latent behavioral patterns that closely align with expert evaluations. By adapting their internal representations over time, RNNs can closely mimic the decision-making processes observed in experimental settings.

RAINSTORM builds upon these recent advancements in deep learning-based analysis by providing an integrated workflow for tracking mice behavior in object recognition tasks. One of its key advantages is the ability to incorporate manual annotations into model training, ensuring flexibility across different experimental conditions. This tool’s multiple labeling approaches—manual, geometric, and neural network-based—allow users to compare methods and validate their results. After analyzing exploratory behavior using the Geometric and/or Automated Behavioral Labeling protocols, the resulting labeled file can be used as input on the Manual Behavioral Labeling protocol to run a frame-by-frame assessment of the obtained labels.

This tool was designed and tuned using behavioral videos of C57BL6 mice performing object recognition tasks. The objects in these experiments measure approximately 2 cm in both height and diameter, ensuring that the trained example models work well under these specific conditions. If researchers wish to analyze exploration of targets with different dimensions, it is recommended to train a new neural network using their own data (see section ‘Create the colabels file’ in Notebook 3a-Create_models). Notably, the pipeline was validated with behavioral videos from a different laboratory, using a different pose estimation mode on an alternative experimental setup, which supports its versatility.

Applications extend beyond exploratory tasks. RAINSTORM can be adapted for assessing social interactions, locomotion patterns, and behavioral phenotyping in neurodegenerative and psychiatric disease models. Its modular structure also allows for integration with other computational tools, making it a valuable resource for researchers in neuroscience, experimental psychology, and behavioral sciences in general. Ultimately, RAINSTORM redefines automated behavioral analysis by delivering a scalable, transparent, and highly customizable framework that accelerates discovery across diverse experimental paradigms.

#### Critical Parameters

Parameters in the params.yaml file:

- *path: Path to the experiment folder containing the pose estimation files*
- *filenames: Pose estimation filenames*
- *software: Software used to generate the pose estimation files (‘DLC’ or ‘SLEAP’)*
- *fps: Video frames per second*
- *bodyparts: Tracked body parts*
- *targets: Exploration targets*
- *prepare_positions: Parameters for processing positions:*
  ○ *confidence: How many std_dev away from the mean the points likelihood can be without being erased (it is similar to asking ‘how good is your tracking?’)*
  ○ *median_filter: Number of frames to use for the median filter (it must be an odd number)*
- *geometric_analysis: Parameters for geometric analysis:*
  ○ *roi_data: Loaded from ROIs.json*
    ▪ *frame_shape: Shape of the video frames ([width, height])*
    ▪ *scale: Scale of the video in px/cm*
    ▪ *areas: Defined ROIs (areas) in the frame*
    ▪ *points: Key points within the frame*
  ○ *distance: Maximum nose-target distance to consider exploration*
  ○ *orientation: Set up orientation analysis*
    ▪ *degree: Maximum head-target orientation angle to consider exploration (in degrees)*
    ▪ *front: Ending bodypart of the orientation line*
    ▪ *pivot: Starting bodypart of the orientation line*
  ○ *freezing_threshold: Movement threshold to consider freezing, computed as the mean std of all body parts over 1 second*
- *automatic_analysis: Parameters for automatic analysis:*
  ○ *model_path: Path to the model file*
  ○ *model_bodyparts: body parts used to train the model*
  ○ *rescaling: Whether to rescale the data*
  ○ *reshaping: Whether to reshape the data (set to True for RNN)*
  ○ *RNN_width: Defines the shape of the Recurrent Neural Network*
    ▪ *past: Number of past frames to include*
    ▪ *future: Number of future frames to include*
    ▪ *broad: Broaden the window by skipping some frames as it strays further from the present*
- *seize_labels: Parameters for the analysis of the experiment results:*
  ○ *groups: Experimental groups you want to compare*
  ○ *trials: If your experiment has multiple trials, list the trial names here*
  ○ *target_roles: Role/novelty of each target in the experiment*
  ○ *label_type: Label type used to measure exploration (geolabels, autolabels, etc)*

Parameters in the modeling.yaml file:

- *path: Path to the models folder*
- *colabels: The colabels file is used to store and organize positions and labels for model training*
  ○ *colabels_path: Path to the colabels folder*
  ○ *labelers: List of labelers on the colabels file (as found in the columns)*
  ○ *target: Name of the target on the colabels file*
- *focus_distance: Window of frames to consider around an exploration event*
- *bodyparts: List of body parts used to train the model*
- *split: Parameters for splitting the data into training, validation, and testing sets*
  ○ *validation: Percentage of the data to use for validation*
  ○ *test: Percentage of the data to use for testing*
- *RNN: Set up the Recurrent Neural Network*
  ○ *width: Defines the shape of the wide model*
    ▪ *past: Number of past frames to include*
    ▪ *future: Number of future frames to include*
    ▪ *broad: Broaden the window by skipping some frames as it strays further from the present*
  ○ *units: Number of neurons on each layer*
  ○ *batch_size: Number of training samples the model processes before updating its weights*
  ○ *epochs: Each epoch is a complete pass through the entire training dataset*
  ○ *lr: Learning rate*

#### Advanced Parameters

The function ‘build_RNN’ on section 3.1.5 builds a Recurrent Neural Network, and takes variables that determine the depth and size of each hidden layer of this Deep Learning mechanism:

- units = [32, 24, 16, 8]: Adding units, regardless their size, would include more hidden layers into the architecture.
- batch_size = 64: The batch size is the number of training samples the model processes before updating its weights.
- lr = 0.0001: The Learning Rate controls how much the model adjusts its parameters on each iteration.
- epochs = 60: Each epoch is a complete pass through the entire training dataset.
- Early stopping: Once the validation loss stops decreasing from epoch to epoch, it is better to stop the training to avoid overfitting the model to the training data.
- Learning rate schedule function: Apply warm-up epochs, and a progressive decrease in the learning rate to fine tune the established connections.
  ○ warmup_epochs = 6: Number of warm-up epochs
  ○ initial_lr = 6e-5: Starting learning rate
  ○ peak_lr = 2e-4: Peak learning rate
  ○ decay_factor = 0.9: Decay factor

## Acknowledgementss

We are grateful to Candela Medina (*Laboratorio de Neurofarmacología de los Procesos de Memoria, Cátedra de Farmacología, Facultad de Farmacia y Bioquímica, Universidad de Buenos Aires*) for her help in validating the pipeline and for her creative input, which greatly contributed to the development of the software.

## Author contributions

Santiago D’hers: Conceptualization, data curation, formal analysis, investigation, methodology, software, validation, visualization, writing-original draft, and editing;

Agustina Denise Robles: Validation, writing-review and editing;

Santiago Ojea Ramos: Validation, writing-review and editing;

Guillermina Bollini: Validation, writing-review and editing;

Mariana Feld: Resources, funding acquisition, supervision, validation, writing-review and editing;

## Data availability statement

All code for the RAINSTORM project can be found at github.com/sdhers/RAINSTORM. All video recordings used to develop and test the protocols are to be made available on demand.

